# A Novel Local Ablation Approach for Explaining Multimodal Classifiers

**DOI:** 10.1101/2021.06.10.447986

**Authors:** Charles A. Ellis, Darwin A. Carbajal, Rongen Zhang, Mohammad S.E. Sendi, Robyn L. Miller, Vince D. Calhoun, May D. Wang

## Abstract

With the growing use of multimodal data for deep learning classification in healthcare research, more studies have begun to present explainability methods for insight into multimodal classifiers. Among these studies, few have utilized local explainability methods, which could provide (1) insight into the classification of each sample and (2) an opportunity to better understand the effects of demographic and clinical variables within datasets (e.g., medication of subjects in electrophysiology data). To the best of our knowledge, this opportunity has not yet been explored within multimodal classification. We present a novel local ablation approach that shows the importance of each modality to the correct classification of each class and explore the effects of demographic and clinical variables upon the classifier. As a use-case, we train a convolutional neural network for automated sleep staging with electroencephalogram (EEG), electrooculogram (EOG), and electromyogram (EMG) data. We find that EEG is the most important modality across most stages, though EOG is particular important for non-rapid eye movement stage 1. Further, we identify significant relationships between the local explanations and subject age, sex, and state of medication which suggest that the classifier learned specific features associated with these variables across multiple modalities and correctly classified samples. Our novel explainability approach has implications for many fields involving multimodal classification. Moreover, our examination of the degree to which demographic and clinical variables may affect classifiers could provide direction for future studies in automated biomarker discovery.

## 1. INTRODUCTION

More studies have begun to integrate multimodal data for classification problems in healthcare research [1]–[3]. The use of complementary modalities offers a key advantage over a single modality by enabling richer feature extraction and improved classification performance [1]. However, the use of multimodal data also undermines the explainability. Many studies have presented explainability approaches for single modality classifiers [4]–[6], but fewer studies have presented approaches for multimodal classifiers [2][3][7].

Explainability methods consist of global and local methods. Global methods explain the importance that each feature generally has to the model, while local methods show the importance of each feature to the classification of each individual sample [8]. Local explanations can also be combined across a dataset to approximate a global explanation [7][9]. Additionally, local explanations can provide insight into how demographic and clinical variables (e.g., subject age and gender) could be learned by the classifier and affect model performance.

Most multimodal explainability studies have presented global explainability approaches [3][10], such as forward feature selection and ablation. However, to the best of our knowledge, only a couple of multimodal studies have utilized local explainability approaches [7][11]. In [7], the authors used layer-wise relevance propagation to gain global insights, and in [11], the authors occluded portions of an input EEG signal to identify time points that were most important to the classification of a sample. However, those studies have not used the approaches to explore patterns learned by the classifier that are not explicitly part of the data.

Here, we present a novel local ablation approach that provides insight into the importance of each modality to the classification of each sample. We also demonstrate how local explanations can be used to identify the effects of demographic and clinical variables upon a classifier. We utilized automated sleep stage classification as a use-case because sleep stages are well-characterized and there are clinical needs for improved explainability in sleep stage classification. We trained a one-dimensional convolutional neural network (1D-CNN) with electroencephalogram (EEG), electrooculogram (EOG), and electromyogram (EMG) data. We then applied our novel local ablation approach and examined the relationships between the local explanations and demographic and clinical data (i.e., age, sex, and medication).

## 2. METHODS

Here, we describe the methods used in our study. We trained a 1D-CNN to classify between sleep stages using multimodal EEG, EOG, and EMG data. We implemented a novel local ablation approach to identify the importance of each modality to the classification of each sample and performed statistical analyses to identify relationships between demographic and clinical variables and the local ablation results.

### 2.1. Description of Data

We analyzed Sleep Telemetry Data from the Sleep-EDF Database [12] on Physionet [13]. The dataset included recordings from 22 subjects including 7 male and 15 female subjects. Subject age had a mean (μ) of 40.18 years and a standard deviation (σ) of 18.09 years. Each subject had two recordings - one after placebo and one after temazepam – for a total of 44, approximately 9-hour recordings. Each recording had 2 EEG electrodes (FPz-Cz and Pz-Oz), 1 EOG electrode, and 1 EMG electrode with sampling frequencies of 100 Hertz (Hz), a polysomnogram of sleep stages, and a marker tracking recording errors. Sleep stages consisted of Awake, Movement, Rapid Eye Movement (REM), Non-REM 1 (NREM1), NREM2, NREM3, and NREM4.

### 2.2. Description of Data Preprocessing

We separated each recording into 30-second samples, discarded movement and recording error samples, and consolidated the NREM3 and NREM4 samples into a single NREM3 class. In each recording, we z-scored each modality independently. Our dataset consisted of 42,218 samples. The dataset was imbalanced with 9.97%, 8.53%, 46.8%, 14.92%, and 19.78% belonging to Awake, NREM1, NREM2, NREM3, and REM, respectively. Of the two available EEG electrodes, we only used Fpz-Cz.

### 2.3. Description of CNN

We used a 1D-CNN architecture originally developed for EEG sleep staging [14]. Further details on the model hyperparameters, architecture, and training approach are detailed in [10]. For each of 10 folds, we randomly distributed different groups of 17, 2, and 3 subjects to training, validation, and test sets, respectively. We used precision, recall, and F1 scores to quantify classification performance.

### 2.4. Description of Local Ablation Approach

To gain insight into the importance of each modality to the classifier, we applied a local ablation approach. For our novel local ablation approach, we (1) identified the classification probability associated with the top predicted class for an unablated sample, (2) replaced one modality with a combination of Gaussian noise and a sinusoid, (3) identified the corresponding classification probability predicted for the ablated sample, (4) calculated the percent change in classification probability between the top class of the unablated sample and the same class for the unperturbed sample, and (5) and repeated for each modality. We repeated this for each sample in test set of each fold. Similar to an existing global ablation approach [10], we used Gaussian noise (μ = 0, σ = 0.1) and a 40 Hz sinusoid with the goal of mimicking the line-related noise that commonly appears in electrophysiology data. While line-related noise typically appears at 60 Hz, a 100 Hz sampling rate will cause a 60 Hz signal to alias to 40 Hz. After outputting the local ablation results for each fold, we visualized the results over time to identify patterns of explainability that corresponded to activity in different modalities and correct or incorrect classification. We also visualized the local results in a global manner to gain better insight into the classifier as a whole. This global visualization showed the local results of all samples across folds for each classification group (ex. REM classified as NREM1 or NREM3 as Awake).

### 2.5. Description of Statistical Analyses

We are interested in understanding whether there was any relationship between the local ablation results and demographic (i.e., age and sex) and clinical (temazepam or placebo) variables. To gain this insight, we looked at the local ablation results for each modality within each correct classification group (15 tests for each demographic or clinical factor). For Age, we calculated the Pearson’s correlation between the age associated with each sample and the absolute values of the local ablation results. For medication, we performed an unpaired, two-tailed t-test between the absolute values of ablation results for samples from placebo versus samples from temazepam recordings. To determine whether subjects’ sex had an effect, we performed a two-tailed t-test between the absolute values of ablation results for samples from recordings of males versus of females.

## 3. RESULTS AND DISCUSSION

Here we describe and discuss the results of our local ablation and statistical analyses.

### 3.1. Classification Results

Table 1 shows the μ and σ of the F1, precision, and recall of the CNN for each sleep stage. NREM2 had the best F1 score, followed by Awake. NREM2 samples composed a large portion of the data, maybe explaining the performance of the classifier for that class. The NREM1 stage had the lowest performance across all performance metrics, likely because it represented such a small portion of the data. While the Awake and NREM1 classes had comparable numbers of samples, the performance for Awake was likely much higher because Awake samples are so distinct from NREM and REM.

**Table 1.** Classification Performance Results

|  | **Awake** | **NREM1** | **NREM2** | **NREM3** | **REM** |
| --- | --- | --- | --- | --- | --- |
| **F1** | 71.25±05.15 | 39.86±07.19 | 73.28±04.76 | 64.15±15.25 | 65.92±06.28 |
| **Precision** | 72.25±07.12 | 36.20±03.98 | 79.35±03.92 | 56.78±18.35 | 69.04±07.14 |
| **Recall** | 70.90±07.02 | 46.28±13.52 | 68.71±08.51 | 78.22±10.24 | 63.26±06.69 |

### 3.2. Local Visualization of Ablation Results

Figure 1 shows the local ablation results for the first 2 hours of a recording from Subject 12. It shows a sleep cycle starting with Awake samples and ending with REM samples. When classifying the Awake stage, ablating EOG resulted in the highest percent change in probability. As such, among the three modalities, the CNN model mostly relied upon EOG when classifying samples as Awake (0 to 20 minutes). Also, EEG played a dominant role for NREM2 and NREM3 classification and a relatively minor role for NREM1 (20 to 100 minutes). It is noteworthy that the percent change increased in successively deeper stages of NREM sleep. For REM classification (100 to 120 minutes), the CNN placed a comparable level of importance upon EEG and EOG earlier during REM when more REM samples were correctly classified. However, deeper into REM, the CNN placed greater importance on EEG relative to EOG and EMG. This may indicate that undo attention to EEG may have resulted in poor classification performance for REM samples. The negative percent change for some samples meant that the model had increased confidence in their classification when a modality was ablated. It should be noted, however, that many such samples were misclassified, so the removal of that modality only made the classifier more confident in an error.

**Figure 1.**
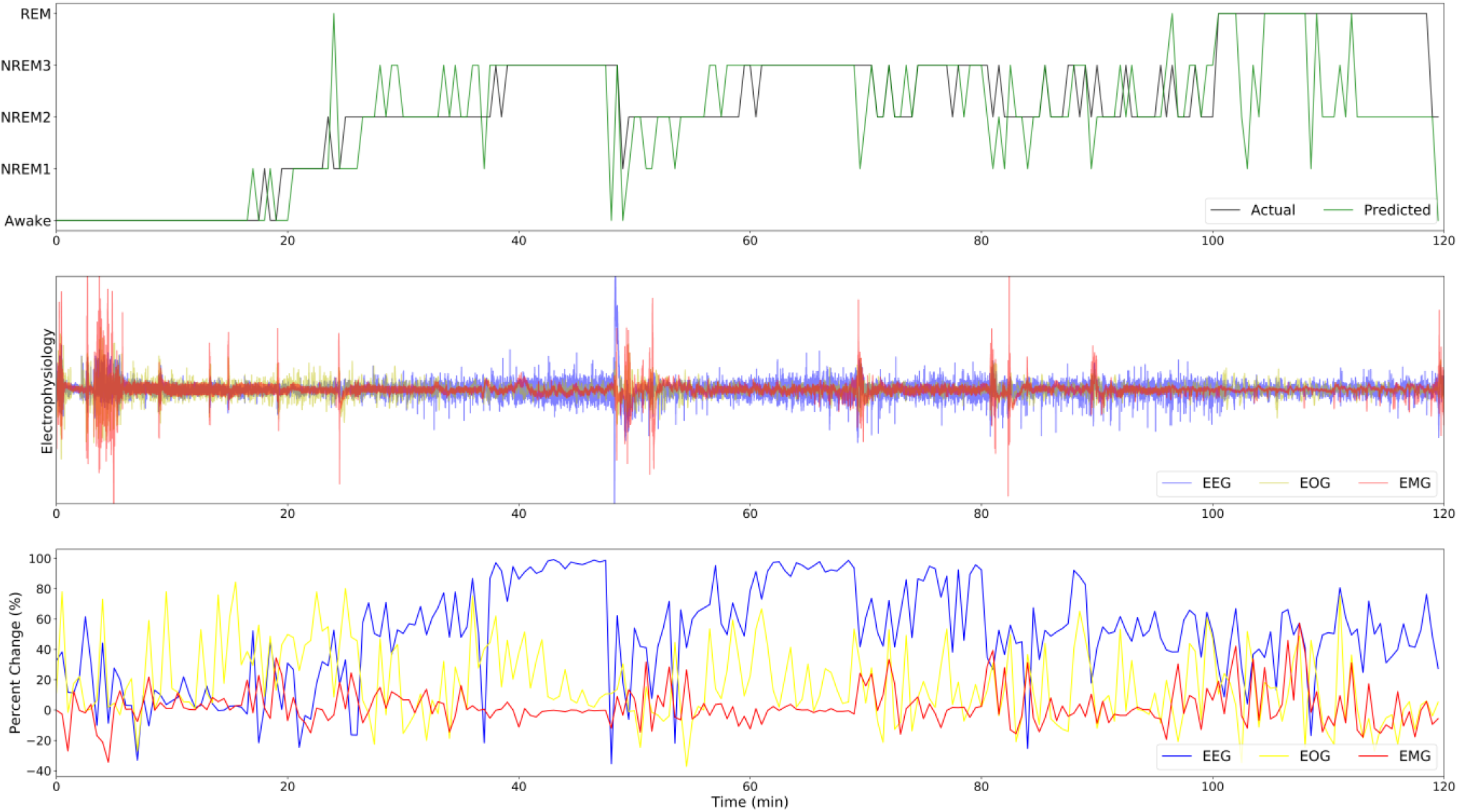
Local Ablation Results for 2 Hours of Recording. The top panel shows the predicted sample classes (green) and the actual sample classes (black). The middle panel shows the data from the different modalities: EEG (blue), EOG (yellow), and EMG (red). The bottom panel shows the importance of each modality to the classification of each sample. The x-axes are aligned and run from 0 to 120 minutes.

### 3.3. Global Visualization of Local Ablation Results

Figure 2 shows the local ablation results in a global manner. In general, EEG seemed to be the most important modality for correctly classified samples. This was clearly the case for correctly classified Awake, NREM2, and NREM3 samples. EEG was also most important for the correct classification of REM, though EOG was nearly as important. Interestingly, EOG was clearly most important to the correct classification of NREM1. This is unexpected, as EOG is typically used for identifying Awake and REM [15]. The reliance of the classifier upon EOG for identifying NREM1 could explain the relatively poor performance of the classifier for NREM1 and could be a factor of the smaller class size. While we can compare the results between modalities within each classification group, it is also worth comparing the importance of each modality across classification groups.

**Figure 2.**
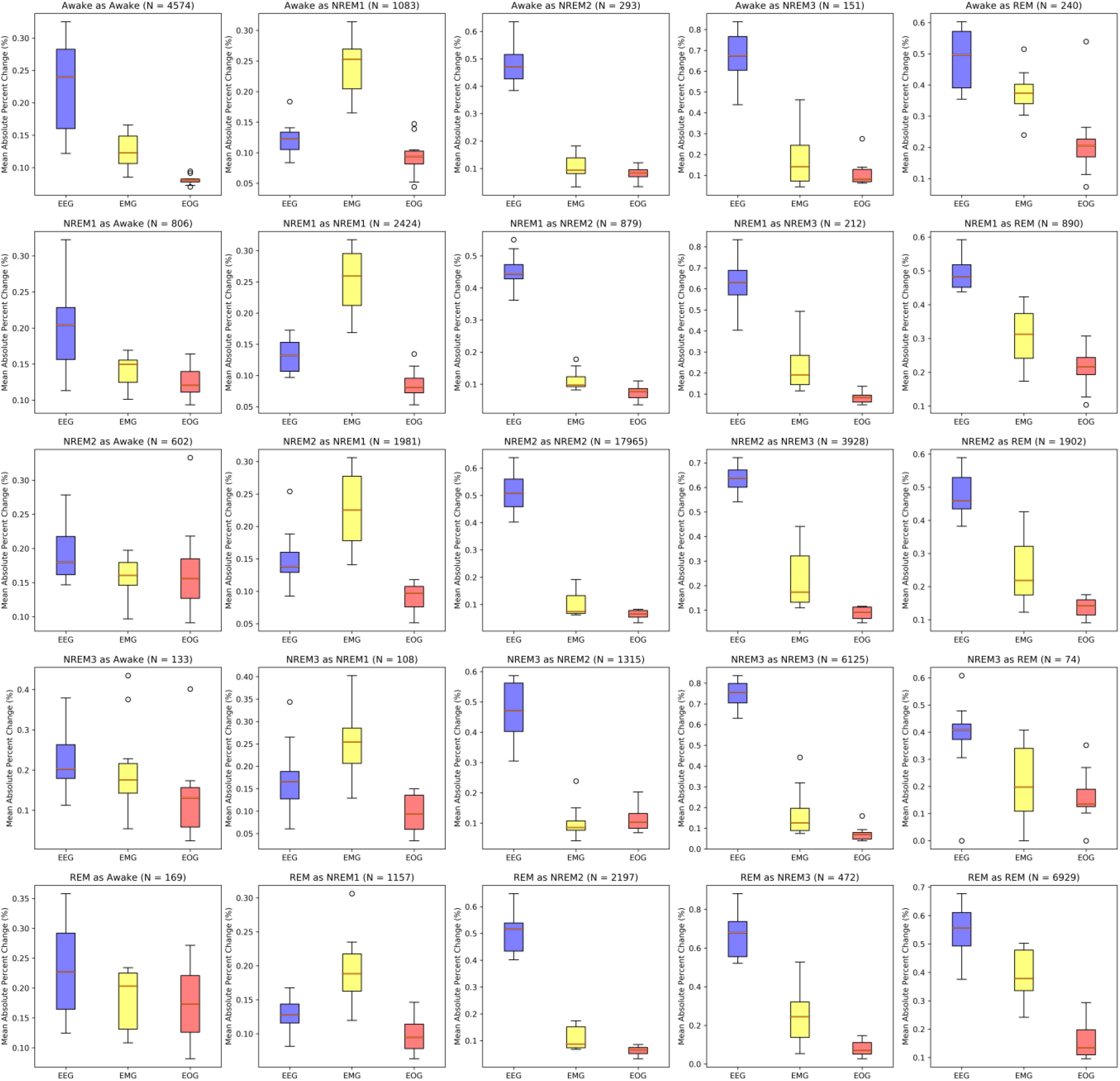
Mean Absolute Percent Change in Classification ProoaDility Across Folds. Each panel shows the μ absolute percent change in top class classification probability for all samples within each fold and classification group. The μ absolute percent change for correctly classified samples within each fold is shown in the panels on the left-to-right diagonal. Off-diagonal show the μ absolute percent change for incorrectly classified samples. EEG, EOG, and EMG correspond to blue, yellow, and red boxes, respectively.

The ablation of modalities in correctly classified NREM2, NREM3, and REM classes resulted in very large importance values (left-to-right diagonal of Figure 2). For EEG, NREM2, NREM3, and REM had median values of around 50%, 73%, and 55%, respectively. In contrast, for correctly classified Awake and NREM1 groups, ablation of EEG had a median change of around 24% and 13%, respectively. Additionally, ablation of EOG had a larger effect upon REM than upon NREM1. While this was the case, EOG was much more important for NREM1 relative to EEG than for REM relative to EEG. For Awake, the classifier obtained performance comparable to or better than the other classes [10]. That fact along with how ablation of each modality seemed to have a smaller effect upon the classification of Awake samples could indicate that the classifier developed a robust representation of the Awake class and could rely upon other modalities when one modality was ablated. It could also indicate that there were features associated with Awake in each of the modalities. This would fit with sleep scoring guidelines, as EEG during Awake tends to be more chaotic than in other sleep stages [15] and EOG and EMG would be expected to have more activity if someone were moving their eyes or face while awake.

Interestingly, for samples incorrectly classified as Awake, the classifier seemed to rely more on EMG and EOG and less on EEG relative to correctly classified Awake samples [here you already do what I suggested above]. This could fit with the possible increased robustness of patterns learned by the classifier for Awake samples. If more patterns were learned, it would be possible for those patterns to affect incorrect classifications. Samples incorrectly classified as NREM1 and NREM2 had similar patterns of importance to those samples that were correctly classified as NREM1 and NREM2. Samples incorrectly classified as NREM3 seemed to have greater variance in EEG and EOG importance across folds. This could indicate that the classifiers relied upon similar patterns to make correct classification across folds and dissimilar patterns for incorrect classification across folds. The dissimilar patterns could be a factor of differences in training datasets. Also, correctly classified NREM3 samples seemed to give greater and lesser importance to EEG and EOG, respectively, relative to samples incorrectly classified as NREM3. Lastly, the classifier seemed to give greater importance to EEG and EOG for samples correctly classified as REM than for samples incorrectly classified as REM. For Awake and NREM1 samples classified as REM, the classifier seemed to prioritize EMG more than it did for correctly classified REM samples.

### 3.4. Statistical Analyses

Figure 3 shows the results of the statistical analyses seeking a relationship between the local ablation results and demographic and clinical information. Interestingly, the local ablation values seemed to have significant relationships with the demographic and clinical variables across most of the modalities and classification groups. If the explanations had no relationship with the demographic information, that would likely mean that there were no patterns learned by the classifier associated with the demographic and clinical variables. The existence of significant relationships in the majority of instances supports the existence of EEG, EMG, and EOG patterns across sleep stages associated with medication, sex, and age. Interestingly, placebo samples had higher EEG importance than temazepam samples across all sleep stages except for NREM2 and REM. For subject sex, males seemed to have greater importance in EOG than females and corresponding lower levels of EMG and EEG importance. Lastly, REM EOG was highly affected by age, maybe indicating changes in REM EOG activity with age.

**Figure 3.**
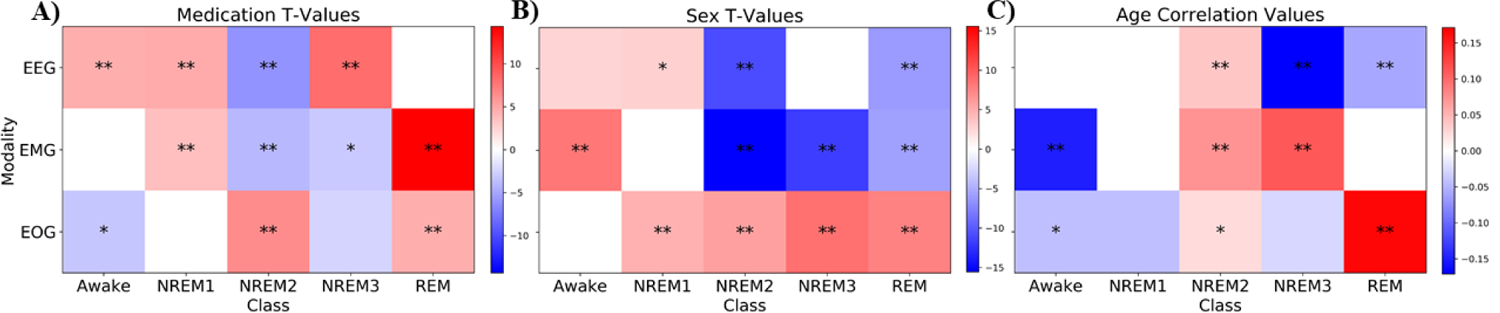
Statistical Analyses of Local Explanations and Demographic and Clinical Variables. Panels A and C show heatmaps of the t-statistics resulting from paired t-tests comparing the local results for all test samples belonging to placebo versus temazepam and male versus female groups, respectively. Panel C shows heatmaps of the coefficients for correlation analyses between the age of the subject associated with each sample and the local ablation results for that classification group. The y-axis indicates the modality of each row (i.e. EEG, EMG, and EOG, from top to bottom). The x-axis indicates the class of each column. Squares that are whited out represent insignificant p-values. Squares with color have significant p-values (p<0.05), and squares that have one or two asterisks have p-values of p < 0.01 or p < 0.001, respectively. For the purpose of interpreting coefficient values, it should be noted that the t-test for medication (Panel A) were performed as Placebo versus Temazepam (i.e., a positive coefficient indicates higher importance for placebo) and that the t-test for sex (Panel B) were performed as Male versus Female (i.e., a positive coefficient indicates higher importance for male).

In most cases, samples that were correctly classified had significant relationships with the local explanations. However, there were several instances in which there were no significant relationships (e.g. NREM1 EMG and NREM2 EOG for medication). This indicates that Medication, Sex, or Age likely do not have an effect upon that particular modality and class, or that the model did not learn any discriminatory patterns associated with the demographic variables for samples belonging to that particular modality and class.

### 3.5. Shortcomings and Next Steps

While our local ablation approach offers features that existing global approaches do not offer, the use of ablation can be problematic. Ablation can potentially create out-of-distribution samples that may lead to an incorrect importance estimates [8]. We ablated modalities with signals mimicking line-related noise in an effort to reduce the likelihood of creating out-of-distribution samples [10]. Nevertheless, other local explainability approaches might provide more reliable explanations [7]. Additionally, it is possible to use global methods to gain insight into the effects of modality ablation upon how samples are both correctly and incorrectly classified by measuring the change in a confusion matrix that occurs following ablation [11]. It would be interesting to compare global results of that nature with the results that we obtained using a local approach in Figure 2.

## 4. CONCLUSION

We presented a novel local explainability approach for insight into classifiers trained on multimodal time-series data with automated sleep stage classification as a use-case. We demonstrated how local methods can be visualized locally for insight into the evolving importance of modalities over time and globally for insight into the correct and incorrect classification of samples. We further demonstrated how local approaches can be used to examine the degree to which demographic and clinical variables may be integrated into classifiers not targeting those characteristics. Moving forward, we hope that our novel explainability approach and variable analysis will enable higher resolution insight into classifiers across a variety of application areas involving multimodal data.

